# Bedrock radioactivity influences the rate and spectrum of mutation

**DOI:** 10.1101/2020.03.23.003616

**Authors:** Nathanaëlle Saclier, Patrick Chardon, Florian Malard, Lara Konecny-Dupré, David Eme, Arnaud Bellec, Vincent Breton, Laurent Duret, Tristan Lefébure, Christophe J. Douady

## Abstract

All organisms on Earth are exposed to low doses of natural radioactivity but some habitats are more radioactive than others. Yet, documenting the influence of natural radioactivity on the evolution of biodiversity is challenging. Here, we addressed whether organisms living in naturally more radioactive habitats accumulate more mutations across generations using 14 species of waterlice living in subterranean habitats with contrasted levels of radioactivity. We found that the mitochondrial and nuclear mutation rates across a waterlouse species’ genome increased on average by 60 and 30%, respectively, when radioactivity increased by a factor of three. We also found a positive correlation between the level of radioactivity and the probability of G to T (and complementary C to A) mutations, a hallmark of oxidative stress. We conclude that even low doses of natural bedrock radioactivity influence the mutation rate through the likely accumulation of oxidative damage, in particular in the mitochondrial genome.

## Introduction

Natural radioactivity is the main natural source of exposure to ionizing radiations on Earth. Natural radioactivity is generated by cosmic radiation or by radionuclides released from the bedrock. While levels of cosmic radiation fluctuates over time due to cosmic events such as supernovae or solar flares, bedrock radioactivity remained mainly stable until 2 billion years ago, when it began to slowly decrease (Karam and Leslie, 2005). Bedrock radioactivity depends on the nature of the rocks which extensively varies spatially (*e.g* Ielsch et al., 2017). While few extremely naturally radioactive sites such as the India Kerala and Iranian Ramsar region have been monitored for their impact on the human mutation rate (Forster et al., 2002; Masoomi et al., 2006) or on plant physiology (Saghirzadeh et al., 2008), the influence of regional variation in baseline natural radioactivity on the evolution of biodiversity is still unknown (Møller and Mousseau, 2012).

Natural radioactivity can modify the rate of molecular evolution by increasing the rate of mutations. Ionizing radiations can damage DNA directly by breaking the DNA sugar-phosphate backbone, or indirectly by the radiolysis of water in cells which results in the production of reactive oxygen species (ROS), which are mutagenic. These impacts have been well characterized after exposure to high doses of radioactivity (Dubrova et al., 1996; Ziegler et al., 1993) but the multi-generational impact of exposure to low doses of radioactivity is poorly known. Some authors (Tubiana et al., 2006, 2009) propose that DNA repair may completely counteract the effect of ionizing radiations for doses below 0.1 Gy, suggesting that low doses have no biological impact. However, *in vitro* exposure to very low dose of ionizing radiation increases the mutation rate of mammalian cells (Vilenchik and Knudson, 2000), suggesting that the repair system is not activated at low radiation doses, thus facilitating the accumulation of mutations that may be transmitted across generations. While some studies found an increase in the number of mutations in the offsprings of exposed people (Dubrova et al., 1996; Forster et al., 2002), others studies found the opposite (Satoh C et al., 1996; Czeizel et al., 1991), leaving open the question of the transmission of mutations generated by low doses of radioactivity. Studying the long-term mutational impact of natural radioactivity is challenging because it raises a number of methodological difficulties. On the one hand, experimentally exposing multiple generations of multicellular organisms to low doses of radiation would require years of experimentation and complex experimental controls. On the other hand, the main obstacles to *in naturae* studies are (i) the organisms’ mobility, which prevents certainty that a population was exposed to the same natural radioactivity for many generations, and (ii) confounding factors such as ultraviolet radiation from the sun. Here we overcome these difficulties by coupling *in situ* radioactivity characterizations with the distinctive bio-ecological characteristics of subterranean waterlice within a phylogenetic comparative framework. Subterranean waterlice are never exposed to UV radiation, live in contrasted bedrock set-ups and have very limited dispersal capacity (Eme et al., 2018), allowing us to make the assumption that different species have persisted in different but nearly constant radioactive habitats for numerous generations.

## Results and Discussion

In order to build a robust and powerful comparative design aimed for testing the influence of natural radioactivity on the mutation rate, we first prospected for closely related subterranean species living in contrasted radioactive set-ups. Using the map of bedrock uranium content in France (Ielsch et al., 2017), we prospected areas with low and high radioactivity. From this large survey (58 sites with waterlice), we selected 14 sites roundwith contrasted levels of *α* radioactivity which were inhabited by closely related groundwater waterlice species. We paid special attention that *α* radioactivity differed at least by a factor of three between two habitats, each containing a closely related species (Figure 1, Table S1). On average low level of radioactivity was around 0.357 Bq/g of dry sediment and high level around 1.259 Bq/g of dry sediment. Based on transcriptome sequencing and *de novo* assembly, we used a phylogenetic approach to estimate nuclear and mitochondrial synonymous substitution rates (*i.e.* the rate of mutations which are fixed). While experimental approaches allow to measure the impact of radioactivity on somatic mutations or on the transmission of mutations across few generations, this phylogenetic approach allows us to measure the impact of natural radiation on the germinal mutation rate over the course of a species history.

**Figure 1:**
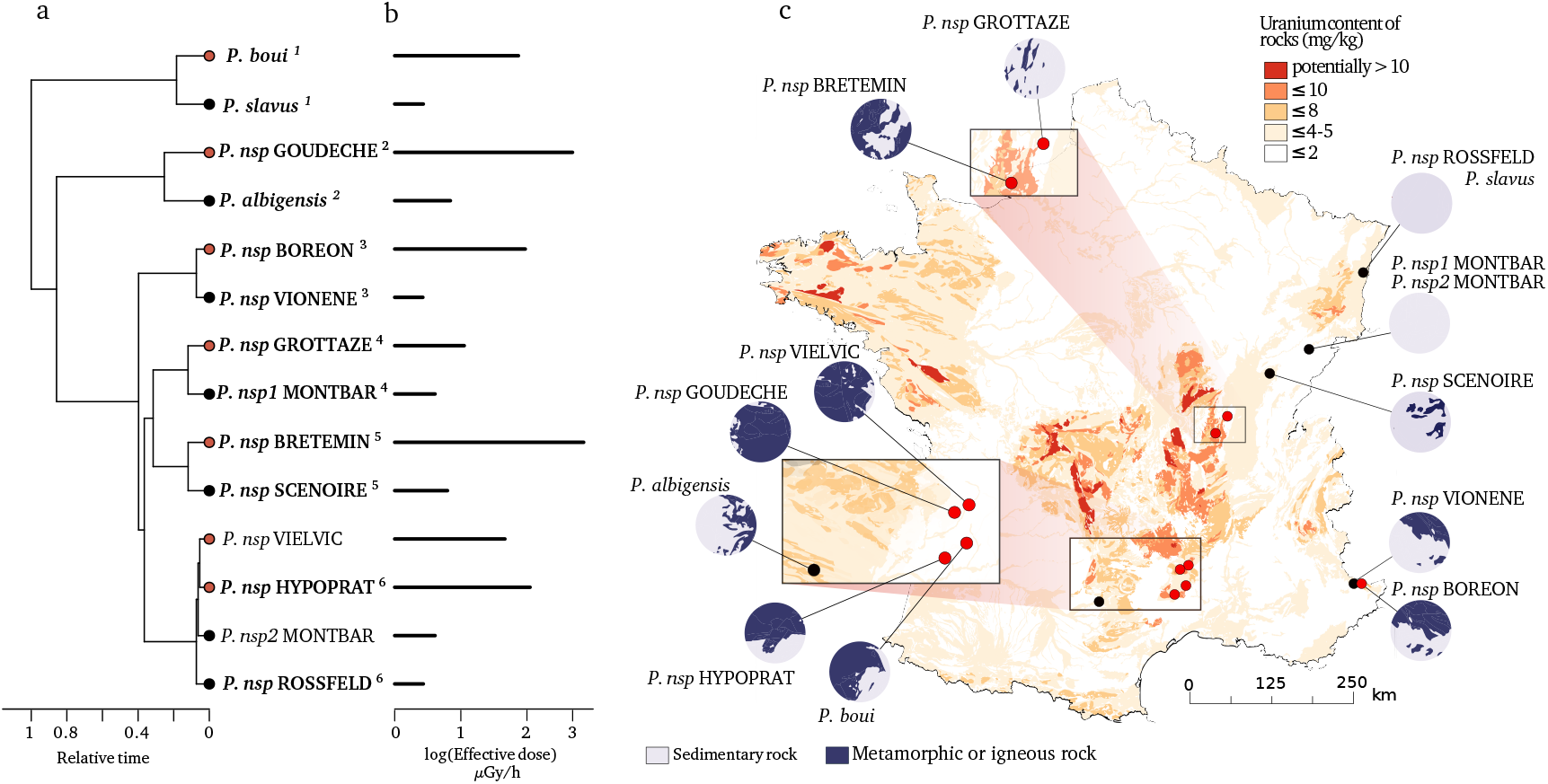
Species and locations selected to study the impact of bedrock radioactivity on the mutation rate and spectrum. 14 species with contrasted bedrock radioactivity exposure were selected (black dot: low exposure, red dot: high exposure). Based on their phylogenetic history (a), we further selected 6 monophyletic pairs of closely related species to compare their mutational spectrum (Vielvic and Montbar are excluded because of unresolved phylogeny, pairs are indicated using superscript numbers). Received dose of radioactivity was measured on sediments of the sampled sites (b). For each site, the areal proportion of low-radioactivity sedimentary rocks and high-radioactivity metamorphic and igneous rocks in a radius of 15 km around the sampling (*λ*15) site is represented with circles next to the map (c).

Using the 14 selected species and locations, we tested whether there was a significant positive relationship between natural radioactivity and the long-term mutation rate. The latter was estimated using the synonymous substitution rate (*d*_S_) calculated on the terminal branches of the phylogenetic tree tracing the history of these 14 species. The *d*_S_ is the rate at which silent mutations accumulate in protein coding genes and, when calculated using many different loci and in the absence of a strong synonymous codon usage bias (see methods), is an estimator of the average mutation rate across a species’ genome (Kimura, 1983). We computed *d*_S_ using 769 one-to-one nuclear and 13 mitochondrial orthologous protein-coding genes shared by all 14 species. At each sampling site, we measured the global *α* radioactivity and the activity of all radio-elements in the sediment. The analysis of the composition in radionuclides at each site reveals that 2 sites (BRETEMIN and BOREON) show a disruption of the secular equilibrium in the U-238 chain. This suggests that nearby industrial activities (*e.g.* lead mines) have modified the natural radioactivity at these two sites. As these industrial activities are very recent (since 1950), their impact on the substitution rate, which is measured on a much longer time scale, is unlikely. We therefore did not use these two sites to test the correlation between *d*_S_ and any site-specific radioactivity measurement (however, see next paragraph for a regional measurement). The *d*_S_ is positively correlated with the alpha radioactivity in the nuclear genome as well as in the mitochondrial genome (Table 1, Figure 2). A linear model predicts a *d*_S_ increase of 31.8% in the nuclear genome and 56.5% in the mitochondrial genome between species living in low (on average 0.357 Bq/g of dry sediment) and high radioactivity (on average 1.259 Bq/g of dry sediment). We also modeled the biologically effective dose of radioactivity received by each species (in *μ*Gy/h, Figure 1). This measure takes into account the transfer coefficient from environment to biota and the radio-toxicity of each radio-element for a crustacean model (ERICA tool V1.2.1 Brown et al., 2016). Again, we found positive correlations between the *d*_S_ and the received dose of radioactivity (Table 1).

**Table 1:** Phylogenetic generalized least square (PGLS) regressions of the nuclear synonymous substitution rate (*d*_S_) and mitochondrial *d*_S_ against the *α* radioactivity measured in the sediments (*α* radio.), the received dose (RD) of radioactivity modeled with the ERICA tool, and the surface of metamorphic and igneous bedrock within a 15 km radius around the sampling sites (*λ*15). *α* radioactivity and RD were log transformed to fit with the linear model assumptions.

| | Nuclear $d_S$ | | | | Mitochondrial $d_S$ | | | | N taxa |
| --- | --- | --- | --- | --- | --- | --- | --- | --- | --- |
| | Slope | L. Ratio | p. value | $R^2$ | Slope | L. Ratio | p. value | $R^2$ | |
| log($\alpha$ radio.) | 0.034 | 5.995 | **0.014** | 0.393 | 0.506 | 7.895 | **0.005** | 0.482 | 12 |
| log(RD) | 0.038 | 6.51 | **0.011** | 0.419 | 0.491 | 5.981 | **0.015** | 0.392 | 12 |
| $\lambda_{15}$ | 0.076 | 9.039 | **0.003** | 0.476 | 1.097 | 11.680 | **0.001** | 0.566 | 14 |

NOTE — Each line corresponds to one likelihood ratio test between the models with and without the given explanatory variable.

**Figure 2:**
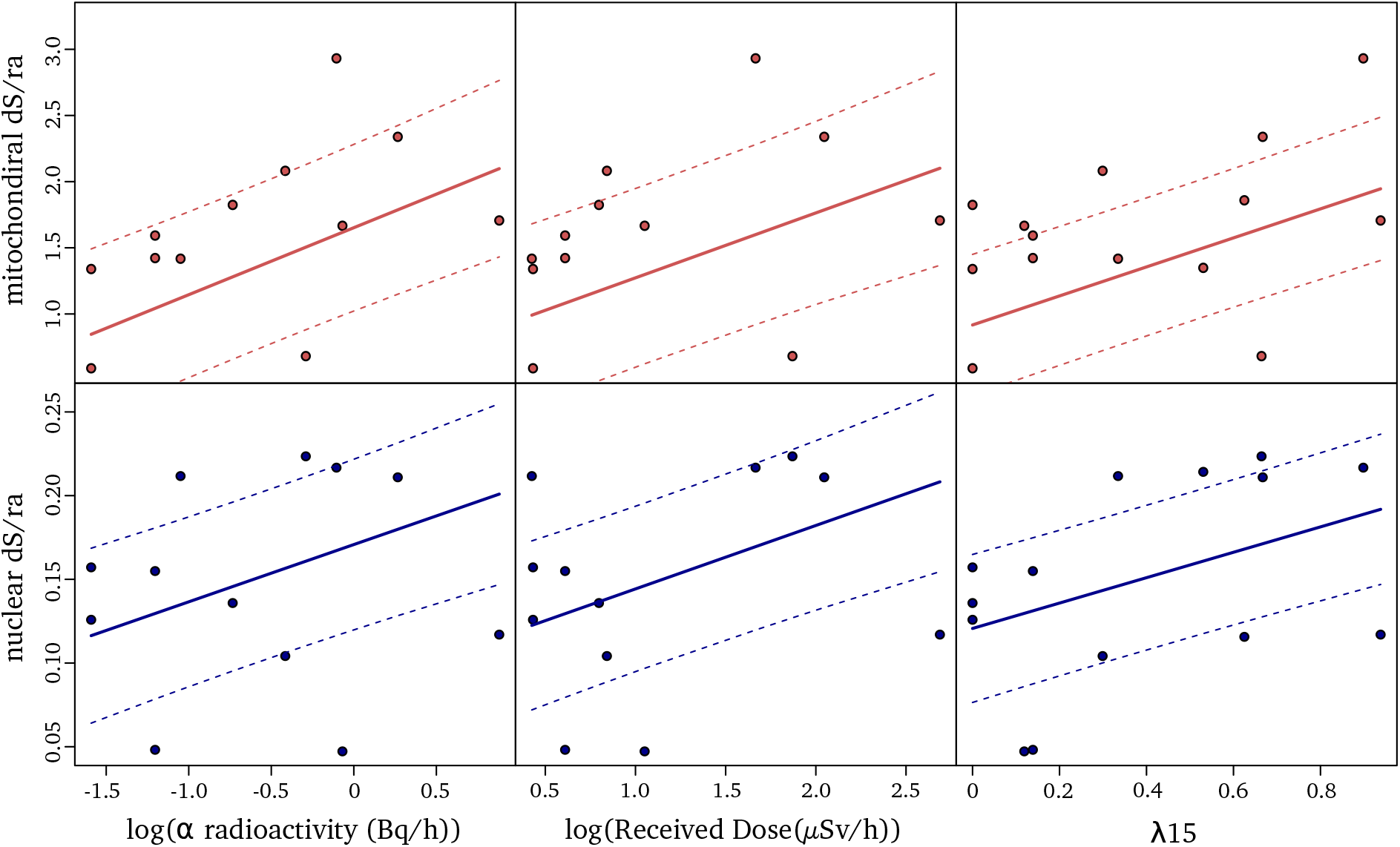
Relationships between synonymous substitution rate relative to the root age (*d*_S_/ra) and radioactivity measured either as the alpha radioactivity measured in sediments (left), the received dose (middle) or the proportion of igneous and metamorphic rock in a radius of 15 km around the sampling sites, *λ*15 (right). Each dot represents a species. *d*_S_ measured on mitochondrial genes are depicted in red and *d*_S_ measured on nuclear genes in blue. The fit of the PGLS model is indicated with a line and the confidence interval of the correlation is indicated with dashed lines.

As previously explained, the measured radioactivity at two sites overestimates the radioactivity level to which the organisms have been exposed for many generations because it is influenced by recent human activities. Moreover, while most species collected in highly radioactive habitats were from metamorphic or igneous formations, two species were from sedimentary formations. Contrary to metamorphic and igneous formations, radioactivity in sedimentary formations is often observed in restricted localites (Ielsch et al., 2017) and can show large variations at the meter scale. A single radiation measurement may not therefore accurately represent the average radiation that a species is exposed to. To account for this variability as well as to include the two human-impacted sites into the regression analysis, we calculated the areal proportion of metamorphic and igneous rock within a 15 km radius around each site (later called *λ*15, Figure 1). This proportion was used as a proxy for the long-term regional radioactive exposure because the average linear distribution range of a groundwater crustacean is 30 km (Eme et al., 2018). We found a positive and stronger correlation between the *d*_S_ and *λ*15 in both genomes (Table 1, Figure 2, n=14 species). The linear model predicts that the nuclear and mitochondrial *d*_S_ of a species living in a metamorphic formation (>50% of metamorphic and igneous rocks) are on average 34.4% and 61.3% higher, respectively, than those of a species living in a sedimentary formation (<50% of metamorphic or igneous rocks).

As bedrock radioactivity is positively correlated with the mutation rate, the underlying question is whether radioactivity also modifies the mutational spectrum, that is, the specific types of mutations that tend to occur. To address this question, we reconstructed the mutational spectrum of 6 independent pairs of species, each composed of two species located in low and high bedrock radiation set-ups, respectively, with a minimum of 3X increase in the received dose of radioactivity between the two species (Figure 1). Briefly, we first estimated species polymorphism across a set of 2490 one-to-one orthologous genes by sequencing transcriptomes for eight individuals per species. After ancestral sequence reconstruction, we then identified mutations that occurred in each species and computed the relative proportion of each type of mutation (from A to T, A to C, …), pooling together complementary mutations (e.g. p(C→A) + p(G→T) = p(C:G→A:T)). We found that bedrock radioactivity was correlated with three types of mutations (Table 2, Figure 3): while the proportions of A:T→T:A and A:T→G:C mutations decreased with increasing radioactivity, the proportion of C:G→A:T mutations increased with bedrock radioactivity, regardless of the radioactivity characterisation (PGLS, p<0.05, Table 2). Selection is unlikely to be responsible for all these correlations as they are also observed when the data set is limited to mutations found at the third, usually redundant, codon position (Figure S2, Table S2). While we found no explanation in the litterature for the decrease of A:T→T:A and A:T→G:C mutations, C:G→A:T mutations, and in particular the G to T mutation triggered by the formation of 8-oxoguanine, are a hallmark of oxidative damage (Shibutani et al., 1991). Similarly, high artificial doses of ionizing radiation were found to increase oxidative damage (Einor et al., 2016; Haghdoost et al., 2006). We thus propose that bedrock radiation impacts the rate and spectrum of mutation of in naturae populations through the formation of reactive oxygen species (ROS, unstable molecules that contain oxygen) generated by the radiolysis of water in cells (Riley, 1994; Azzam et al., 2012).

**Table 2:** Phylogenetic Generalized Least Square (PGLS) regressions of the proportion of each type of mutation against the *α* radioactivity measured in sediment (*α* radio.), the Received Dose (RD) modeled with ERICA tool, and the areal proportion of metamorphic and igneous rock within a 15 km radius (*λ*15). *α* radioactivity and RD were log transformed to fit with linear model assumptions. *R*^2^ are Cox-Snell pseudo *R*^2^.

| Response variable | Explanatory variable | Slope | L.Ratio | P.value | $R^2$ | N |
| --- | --- | --- | --- | --- | --- | --- |
| P(A:T>T:A) | log($\alpha$ radio) | -0.009 | 6.8187 | 0.009 | 0.433 | 12 |
|  | log(RD) | -0.011 | 9.3479 | 0.002 | 0.541 |  |
| | $\lambda 15$ | -0.025 | 6.994 | 0.008 | 0.442 | 14 |
| P(A:T>C:G) | log($\alpha$ radio) | 0.001 | 0.121 | 0.728 | 0.010 | 12 |
|  | log(RD) | 0.000 | 0.015 | 0.904 | 0.001 |  |
| | $\lambda 15$ | 0.001 | 0.005 | 0.945 | 0.000 | 14 |
| P(A:T>G:C) | log($\alpha$ radio) | -0.022 | 7.778 | 0.005 | 0.477 | 12 |
|  | log(RD) | -0.019 | 4.004 | 0.045 | 0.284 |  |
| | $\lambda 15$ | -0.014 | 0.394 | 0.530 | 0.032 | 14 |
| P(C:G>G:C) | log($\alpha$ radio) | 0.005 | 2.137 | 0.144 | 0.163 | 12 |
|  | log(RD) | 0.006 | 2.403 | 0.121 | 0.181 |  |
| | $\lambda 15$ | 0.006 | 0.360 | 0.549 | 0.030 | 14 |
| P(C:G>A:T) | log($\alpha$ radio) | 0.013 | 13.010 | 0.000 | 0.662 | 12 |
|  | log(RD) | 0.014 | 12.079 | 0.001 | 0.635 |  |
| | $\lambda 15$ | 0.042 | 13.791 | 0.000 | 0.683 | 14 |
| P(C:G>T:A) | log($\alpha$ radio) | 0.012 | 3.323 | 0.068 | 0.242 | 12 |
|  | log(RD) | 0.010 | 1.684 | 0.194 | 0.131 |  |
| | $\lambda 15$ | -0.009 | 0.183 | 0.669 | 0.015 | 14 |

**Figure 3:**
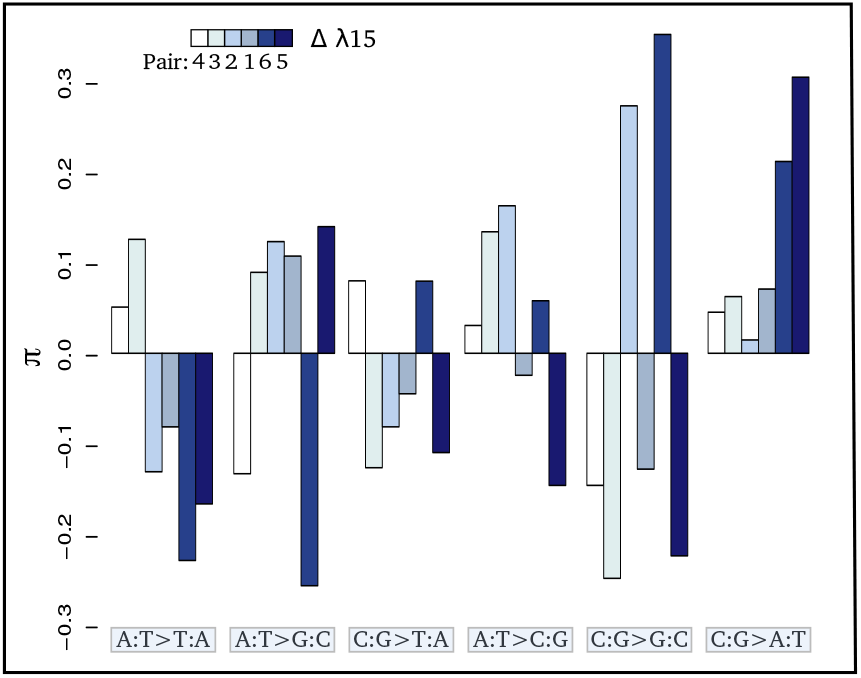
Contrasts (*π*) of the relative proportion of each mutation [p(i:j → k:l)] in each pair of sister species: 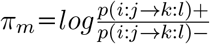 where + and - refer respectively to the species exposed to the higher and lower level of radioactivity in the pair m. Thus, positive bars represent a higher proportion of the given mutation in the species living in the high radioactivity rock. From left to right, bars are in increasing order of difference (Δ) in *λ*15 (the areal proportion of igneous and metamorphic rock in a radius of 15 km around the site) between the two species of each pair. From left to right, mutations are in increasing order of correlation with radioactivity. Numbers below the color scale indicate the species pair number as in Figure 1.

Radioactive environments can cause oxidative stress in two intertwined ways. First, the ionisation of molecules in the cells can directly affect the DNA structure by breaking sugar phosphate backbones or can affect DNA indirectly through the radiolysis of water which decomposes the *H*_2_*O* molecules and create free radicals (Desouky et al., 2015). These free radicals can damage DNA molecules and create mutations. Second, radioactive decay chains also generate heavy metals (lead, polonium,etc) which are toxic for cells and also cause oxidative stress (Quinlan et al., 1988; Pinto et al., 2003). Due to the physicochemical association between radioactivity and heavy metals, *in naturae* experiments cannot discriminate heavy metal chemical toxicity from the direct toxicity of radioactive rays. Therefore, this suggests that naturally radioactive environments impact the rate and spectrum of mutation by directly or indirectly increasing the level of oxidative stress.

Radiolysis of water alone cannot explain the much higher impact of radioactivity on the mitochondrial mutation rate compared to the nuclear rate. Since there is no reason to argue that radiolysis would not evenly occur within cells, it should impact both genomes similarly. However, differences between the two genomes may explain why the mitochondrial genome is more sensitive to radioactivity. First, the mitochondrial genome lacks some repair systems. For example, in a different yet analogous context, UV damage accumulates in the mitochondrial genome while they are repaired in the nuclear genome (Clayton et al., 1974). Second, the two genomes have very different organisations: while the nuclear genome is compacted into chromatin, the mitochondrial one is organized into nucleoids (Chen and Butow, 2005). While the role of the mitochondrial nucleoids is unclear, the chromatin structure and histone proteins protect the nuclear genome against radiation-induced strand breaks (Ljungman, 1991) and oxidative damage (Ljungman and Hanawalt, 1992). Third, direct radioactivity damage to the mitochondria may increase the activity of the mitochondrial respiratory chain and indirectly increase the production of ROS and DNA damage (Yamamori et al., 2012; Kam and Banati, 2013).

While life history traits are known to be central in controlling the mutation rate in metazoans (Nabholz et al., 2008; Martin and Palumbi, 1993), we show here that natural variation of radioactivity can have a comparable effect. Indeed, we found a minimum increase of around 30% percent of the nuclear mutation rate (60% in mitochondria) for species of waterlice living in the more naturally radioactive habitats made of igneous and metamorphic rocks. This increase is of the same magnitude as that observed when waterlice species evolve a 5-fold increase in generation time, a key life history trait controlling mutation rate in waterlice and metazoans in general (Saclier et al., 2018). As groundwater waterlice ingest sediment (Francois et al., 2016), they are internally exposed to radioactivity, which may cause more mutations than through external exposure only (Sawada, 2007). The influence of environmental radioactivity on mutation rate should therefore be explored across a wider range of organisms with contrasted diets.

Natural radioactivity is often considered to have a negligible biological impact (Tubiana et al., 2006, 2009). Indeed, only a handful of isolated studies support an impact of natural radioactivity on the mutation rate. For instance, a higher mutation rate was observed in the human mitochondrial genome in the Kerala region (Forster et al., 2002) and in satellite sequences of crickets inhabiting cave with high radon concentration (Allegrucci et al., 2015). In this study, by combining a large number of genes with the characteristics of the subterranean waterlice, namely the absence of UV confounding effect and limited dispersal, within a statistically powerful comparative framework allowing to work on large time scales and with numerous replicates, we found that a mild variation (*≃* 3.5X) in natural bedrock radioactivity substantially alters the mutation rate, in particular the mitochondrial one. While the universality of this finding warrants corroborative studies in other taxa, it suggests that the influence of natural radioactivity on the evolution of biodiversity may have been overlooked.

## Methods

### Sampling

For 58 sites in France selected on the map of uranium (Ielsch et al., 2017), we collected Asellidae species and sampled about 50 grams of sediment to measure global *α* radioactivity (see the following paragraph). Animals and sediments were collected using the Bou-Rouch pumping methods (Bou and Rouch, 1967). Collected species were stored in 96% ethanol at −20°C and were morphologically and molecularly identified. For molecular identification, DNA was extracted using an optimized chloroform DNA extraction protocol for the Aselloidea (Calvignac et al., 2011). We amplified DNA with primers targeting the 16S mitochondrial rDNA gene. PCR reactions were done following Morvan et al. (2013). PCR products were sequenced in both directions using the same primers as for amplification (GATC Biotech, Konstanz; Eurofins MWG Operon, Ebersberg; SeqLab, Göttingen, Germany; BIOFIDAL, Vaulx-en-Velin, France). Chromatograms were visualized and cleaned using Finch v1.5.0 (Geospiza, Seattle, USA). 16S have been deposited on the European Nucleotide Archive and are available under the accession number from LR214526 to LR214880. Using Eme et al. (2018) molecular species delimitation, each sequence has been assigned to a species. Based on this taxonomic assignment and radioactivity measurement, 14 species were retained for further analyses (Table S1). For these 14 selected species, during a new sampling trip, individuals were flash frozen alive in the field.

### Measures of radioactivity

#### α radioactivity

In order to estimate the global radioactivity in sediments, we measured the *α* radioactivity. An *α* decay occurs when an atom disintegrates by ejecting an *α* particle, *i.e.* a particle made of two neutrons and two protons. The *α* radioactivity should be correlated with the global radioactivity in natural systems. For the 58 prospected sites, 3 samples of about 50 grams of sediment were collected in polyethylene bottles. *α* radioactivity measurements were made by the LABRADOR service (Institut de Physique Nucléaire de Lyon, France) on proportional counter with the NF ISO 18589-6 standard (Data available on Zenodo, (DOI: 10.5281/zen-odo.3356835).

#### Received dose

In order to estimate the received dose of radiation that is impacting organisms, we collected three samples of 100g of fine sediments (<100*μ*m) in each of the selected sites. These sediments were prepared with the NF EN ISO 18589-2 standard and measured by gamma spectrometry in conformity with the NF EN ISO 18589-3 standard using the PRISNA-P analysis platform at the Centre d’Etude Nucléaire de Bordeaux Gradignan (CENBG). This platform is certified by the French Nuclear Security Authority (ASN) for measures of natural radioactivity. Samples were dried in open air, and then dried at 100°C. Matters were packed in a waterproof geometry. Geometries were sealed for one month and then counted for a duration of 86500 seconds on the same chain of measure. The chain used is an ORTEC chain, presenting an efficiency of about 60% and calibrated in May 2016. This chain is equipped with a cosmic veto device and located in a half buried laboratory in order to: (i) attenuate the background noise, (ii) improve the detection limits and (iii) reduce the measure uncertainty. The activity of the main radionuclides were measured in sediment and the activity of the remaining radionuclides was deduced based on the hypothesis of a secular equilibrium of the uranium 238 and thorium 235 chains. As activities of the radionuclides of the uranium 235 decay chain are generally low, only measures higher than the decision threshold (according the measure variability) were taken into account. When the uranium 235 activity was too low to be measured it has been deduced from the uranium 238 activity, using the natural isotopic ratio of 21.6.

The received dose impacting organisms was estimated using the ERICA tool (V1.2.1, Brown et al., 2016) with a ‘crustacean’ model. We assumed that organisms stay 10% of their time on the surface of sediment and 90% inside sediment. All radionuclides available in the tool were taken into account (*i.e U* ^238^, *Th*^234^, *U* ^234^, *Th*^230^, *Ra*^226^, *Pb*^210^, *Po*^210^,*U* ^235^, *Th*^231^, *Pa*^231^, *Th*^227^, *Th*^232^, *Ra*^228^ and *Th*^228^). We used the distribution coefficients proposed by the ERICA tool. Concentration factors proposed by the tool were used when available. If not, we used the concentration factor of the closest biogeochemical element available.

Two sites (BRETEMIN and BOREON) show a disruption of the secular equilibrium in the *U* ^238^ chain. This suggests that nearby industrial activities (*e.g.* lead mines) have modified the natural radioactivity of these two sites. As these industrial activities are very recent (since 1950), their impact on the substitution rate which is measured on a much longer time scale is unlikely. These two sites were removed from the correlation between *d*_S_ and radioactivity measured with the global *α* radioactivity or with the received dose.

#### Proportion of magmatic and igneous rocks in a 15 km radius

Usign the geological map of France (scale: 1/1 000 000, ©BRGM), the areal proportions of magmatic and igneous rocks in a radius of 15 km around sampling sites were computed (noted *λ*15), 30 km represent the distribution range for a subterranean isopod (Eme et al., 2018).

### Transcriptome Sequencing and Assembly

#### Sequencing

For each species, we sequenced transcriptomes from 8 individuals. For each individual total RNA was isolated using TRI Reagent (Molecular Research Center). Extraction quality was checked on a BioAnalyser RNA chip (Agilent Technologies) and RNA concentrations were estimated using a Qubit fluorometer (Life Technologies). Prior to any additional analysis, species identification was corroborated for each individual by sequencing a fragment of the 16S gene. Illumina libraries were then prepared using the TruSeq™ RNA Sample Prep Kit v2 (Illumina). For each species one library was paired-end sequenced using 100 cycles, and the 7 other libraries were single-end sequenced using 50 cycles on a HiSeq2500 sequencer (Illumina) at the IGBMC GenomEast Platform (Illkirch, France). We obtained around 30 million single-end reads per individual and 118 million paired-end reads per species.

#### Assembly

Adapters were clipped from the sequences, low quality read ends were trimmed (phred score < 30) and low quality reads were discarded (mean phred score < 25 or if remaining length < 19 bp) using fastq-mcf of the ea-utils package (Aronesty, 2013). Paired-end transcriptomes were *de novo* assembled using Trinity v2.3.2 (Grabherr et al., 2011). Open reading frames (ORFs) were identified with TransDecoder (http://transdecoder.sourceforge.net). For each assembled component, only the most express ORF was retained.

### Families of Orthologous Genes

Gene families were delimited using an all-against-all BLASTP (Altschul et al., 1990) and SiLix (Miele et al., 2011) on the ORFs delimited in the previous step. We then kept gene families containing the 14 species, with only one sequence for each species in order to remove paralogs. We obtained 2490 families hereafter considered as one-to-one orthologous genes. These genes were aligned with PRANK (Löytynoja and Goldman, 2008) using a codon model and sites ambiguously aligned were removed with Gblocks (Castresana, 2000).

### Species Tree and Gene Trees

The 2490 genes were concatenated and a phylogenetic tree (hereafter called the concatenated tree) was built using PhyML v3.0 (Guindon et al., 2010) under a GTR+G+I model with 100 bootstrap replicates and was rooted using the Slavus lineage (*Proasellus boui* and *Proasellus slavus*) as an outgroup (Morvan et al., 2013). Most nodes have a bootstrap value of 100% (Figure S1). Two nodes have values at 84 and 98% in the clade containing P. nsp VIELVIC; P. nsp HYPOPRAT; P. nsp MONTBAR and P. nsp ROSSFELD. To check the relationship between these 4 species, we built 2490 individual gene trees with PhyML v3.0 under a GTR+G+I model with 100 bootstrap replicates. 29 gene trees strongly support (bootstraps > 90%) the phylogeny of the concatenated tree for this clade, 208 support other various topologies and the remaining 2253 gene trees do not support any relationship in particular for this clade (bootstraps < 90%). Thus, the phylogeny for this clade remained unresolved, possibly as the consequence of a concomitant speciation process of these 4 species. For approaches with pairs of sister species, as we were unable to resolve the phylogeny for this clade, we selected the species living in the highest level of radioactivity (*P. nsp* HYPOPRAT) and the species living in the lowest level of radioactivity (*P. nsp* MONTBAR) among these 4 species to build a pair, resulting in a total of 6 pairs of sister species (*sensu* Felsenstein, 2004).

### Mitochondrial genes

Mitochondrial genes were not present amongst the 2490 genes obtained above. Indeed, owing to a different genetic code in invertebrate mitochondria, mitochondrial ORFs were systematically missed by the ORF caller (Transdecoder). We reconstructed mitochondrial genomes using the *de novo* transcriptome assemblies. Large mitochondrial contigs were built with MITObim (Hahn et al., 2013) by using RNA-seq reads. These contigs were mapped on the assembled mitochondrial genome from the closest possible species (taken from Saclier et al., 2018), allowing us to assemble them. Mitochondrial genomes were annotated using the MITOS web server (Bernt et al., 2013). We recovered the 13 mitochondrial protein-coding genes. Mitochondrial genes were aligned with PRANK (Löytynoja and Goldman, 2008) and sites ambiguously aligned were removed with Gblocks (Castresana, 2000).

### Rate of molecular evolution

We used the synonymous substitution rate (*d*_S_) computed on the terminal branches of the tree as a proxy for the long-term species mutation rate (Kimura, 1983). This proxy is valid in absence of selection on codon usage. To check for the absence of biased codon usage, we computed the effective number of codons on the 2490 orthologous genes (ENC, Wright, 1990)). This number varies between 20 (only a single codon is used for each amino acid) and 61 (all synonymous codons are used with equal frequency for each amino-acid). ENC ranged between 49.17 and 50.48 (Table S1), indicating a moderate codon usage bias, more importantly, they do not correlate with alpha radioactivity (PGLS, p.value = 0.6378). Altogether, the *d*_S_ estimation doesn’t seem impacted by a strongly biased or variable codon usage.

To compute *d*_S_ we first removed some genes showing a conflicting phylogeny. Including genes supporting different phylogenies in a concatenation amounts to constrain a wrong phylogeny for these genes which may biases *d*_S_ estimations. Indeed imposing a wrong gene tree will tend to generate convergent mutations in terminal branches of the tree. To avoid such bias in our *d*_S_ estimation we used ProfileNJ (Noutahi et al., 2016) with a bootstrap threshold of 90% to compute a cost of reconciliation between the concatenated tree and the gene trees. We kept the gene families with a cost of reconciliation of zero and with sequences long enough for all species (at least a half of the alignment) and removed all other genes, resulting in a set of 769 gene families. *d*_S_ were estimated using CoEvol (Lartillot and Poujol, 2011). This software program implements a Muse and Gaut codon model (Muse and Gaut, 1994), with Brownian variation in *d*_S_ and *d*_N_/*d*_S_ along the tree. Bayesian inference and reconstruction of the history of variation in *d*_S_ and *d*_N_/*d*_S_ along the tree is conducted by Markov Chain Monte Carlo (MCMC). Two independent chains were run, and were stopped after checking for convergence by eye and with the tracecomp program included in the Coevol package (effective sample size > 200 and discrepancy between chains < 0.3). Chains were stopped after 7117 generations (4200 generations excluded as burn-in). The age of the root was arbitrarily set to 1, resulting in synonymous substitution rate estimates that are relative to the root age (*d*_S_/ra) (Table S1). In order to ensure that assumptions made by CoEvol on the *d*_S_ evolution along branches don’t bias the *d*_S_ estimation, *d*_S_ were also computed with CodeML (Yang, 2007) using a free ratio model and with the Bio++ suite (Dutheil and Boussau, 2008). For the last one, a non-homogeneous model (NY98 model) was first applied to the alignment with BppML and then the MapNH program (Version 1.1.1) of the TestNH package (Guéguen and Duret, 2018) was used to reconstruct the ancestral states to estimate the number of synonymous substitutions on each branch. CoEvol *d*_S_ was highly correlated with CodeML *d*_S_ (*R*^2^=0.81) as well as with Mapnh *d*_S_ (*R*^2^=0.82). Regarding the correlation with radioactivity, by dividing the CodeML *d*_S_ and the Mapnh *d*_S_ by the divergence time estimated by CoEvol in order to obtained comparable *d*_S_ among all species, we obtained similar results whatever the method used to compute the *d*_S_ (Table S3).

### Mutational spectrum

To compute the mutational spectrum, we used an approach by pairs of sister species. We determined the polymorphism at the population level for each species by mapping the 7 single-end transcriptomes on the assembled paired-end transcriptome with BWA (Li and Durbin, 2009). BAM files were produced with SAMtools (Li et al., 2009), and reads2snps (Gayral et al., 2013) was used to detect polymorphic sites. We then conserved only the 2490 orthologous genes shared by all species to compute the mutational spectrum on the same set of genes.

For the two species of a pair, we reconstructed the ancestral sequence using a parsimonious approach. Namely, for each site in the alignment, if the two species had a single shared allele, this allele was considered as ancestral and the other alleles, if they existed, were considered as derived from the ancestral allele. For each species, we estimated the probability of a mutation in their population, *p*(*i → j|f* (*i*)_*anc*_), by counting each type of mutation, either on all positions (Table S8) or on third positions, corrected by the ancestral base frequency:

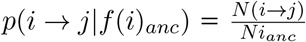

This probability being dependent on the mutation rate *μ*, we estimated the mutational spectrum by the proportion, when a mutation occurs, of mutation from the base i to the base j, noted *p*(*i → j|μ, f* (*i*)_*anc*_):

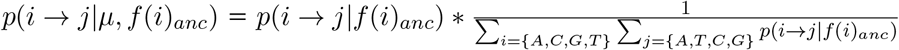

This proportion takes into account the mutation rate and is so comparable across species. We pooled complementary mutations (*e.g.* A to C with T to G) to increase the counts by mutational categories and improve statistical power.

### Statistical analyses

Correlations between *d*_S_ computed on terminal branch of the tree and the different measures of radioactivity were tested using Phylogenetic Generalised Least Squares models (pGLS Martins and Hansen, 1997) with the nlme (Pinheiro et al., 2007) and ape packages (Paradis et al., 2004) in R (R Core Team, 2014). For the *α* radioactivity and effective dose, the two species showing a disruption in the secular equilibrium were removed. The ultrametric tree built by CoEvol was used to calculate the phylogenetic variance/covariance matrix under a Brownian motion model to take into account the non-independence among species in the PGLS. The same statistical procedure was used to test the correlation between the probability of each type of mutation and the effective dose of radioactivity or proportion of metamorphic or igneous rocks, this time pruning P. nsp VIELVIC and P. nsp MONTBAR from the chronogram built by CoEvol. Normality of residuals was checked for all models, log transformation was applied when the normality was rejected (Shapiro test).

## Data availability

16S have been deposited on the European Nucleotide Archive and are available under the accession numbers from LR214526 to LR214880 (https://www.ebi.ac.uk/ena/data/view/LR214526-LR214880).

The 2490 alignments and the list of the 769 genes used to compute synonymous substitution rate have been deposited on Zenodo (https://zenodo.org/deposit/2563829).

Transcriptome reads have been deposited on the European Nucleotide Archive and are available under accession numbers from LR536601 to LR536626 in the study ID PRJEB14193 (https://www.ebi.ac.uk/ena/data/search?query=PRJEB14193). Number of reads and data used for correlations, namely measures of radionuclides and mutations counts have been deposited on Zenodo (https://doi.org/10.5281/zenodo.3356835).

## Aknowledgments

This work was supported by the French program STYGOMICS (CNRS Défi Enviromics), the Zone Atelier Territoire Uranifère, the Grottes d’Azé, the Conseil Départemental de Saône-et-Loire, the Association Culturelle du Site d’Azé and the Agence Nationale de la Recherche (ANR-15-CE32-0005 Convergenomix, France, ANR-17-EURE-0018 H2O’Lyon, France). We gratefully acknowledge support from the CNRS/IN2P3 Computing Center (Lyon/Villeurbanne, France) for providing a significant amount of the computing resources needed for this work. We thank the Grottes d’Azé and more specifically Lionel Barriquand, the Mercantour national parc (authorization n°2015-251) and more specifically its scientific manager Marie-France Leccia, the Fédération Rhône-Alpes de Protection de la Nature and the owner of Bout du monde mine for giving us access for sampling. We thank Marcel Meysonnier, Aymeric and Audric Berjoan, Josiane and Bernard Lips, Audrey Brechet, Claude Bou, Benjamin Benti and Léa Dantony for their help in the field. We are grateful to Nicolas Lartillot, Gilles Escarguel, Marie Sémon, Laurent Guéguen, Nicolas Galtier, Benoît Nabholz and Bastien Boussau for helpful discussions. We also thank Laurent Simon and Laura Grice for their suggestions in the latter stages of manuscript preparation.

## Contributions

Douady C., Lefébure T., Chardon P. and Malard F. designed the study. Saclier N. and Lefébure T. wrote the manuscript. Douady C., Lefébure T., Chardon P., Malard F., Saclier N. and Eme D. did the field work (animal and sediment sampling). Malard F. and Eme D. dissected and identified morphologically all sampled species. Lara Konecny-Dupré extracted DNA and RNA from all samples, made PCR and migrations and prepared library for sequencing. Saclier N. built the phylogeny, computed substitution rate and performed statistical analyses. Duret L. and Saclier N. computed the mutational spectrum. Chardon P. and Breton V. performed all radioactivity measurements and the effective dose estimation. Bellec A. extracted the outcrop cover of low-radioactivity sedimentary rocks and high-radioactivity metamorphic and igneous rocks in a radius of 15 km around the sampling and made the map of uranium for the figure 1. All authors interpreted the data and contributed to the final manuscript.

## Supplementary information

**Figure S1:**
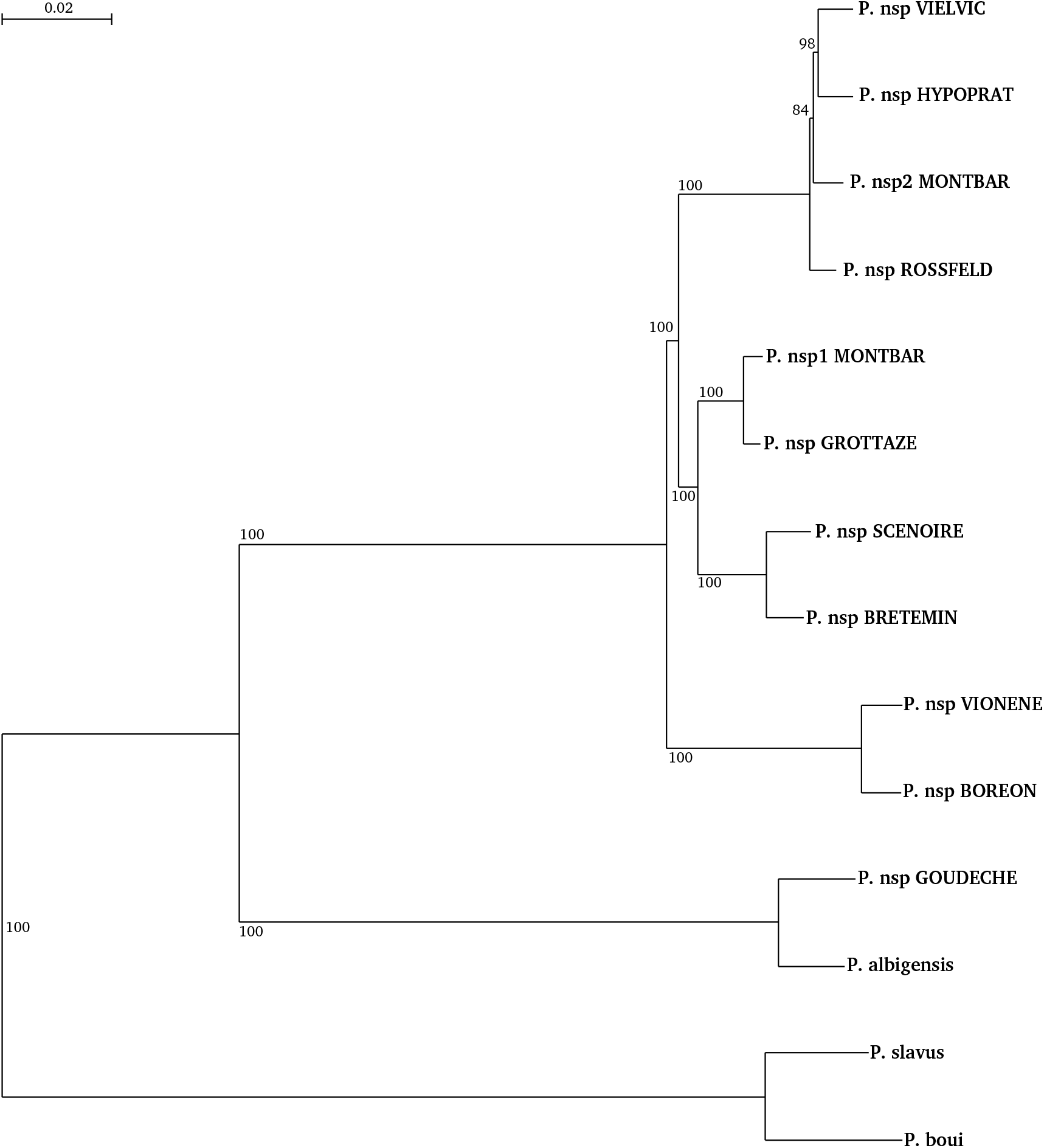
Phylogeny of the 14 species used to compute synonymous substitution rates. Tree was built with the concatenation of the 2490 genes with PhyML3.0, under a GTR+G+I model with 100 bootstrap replicates.

**Figure S2:**
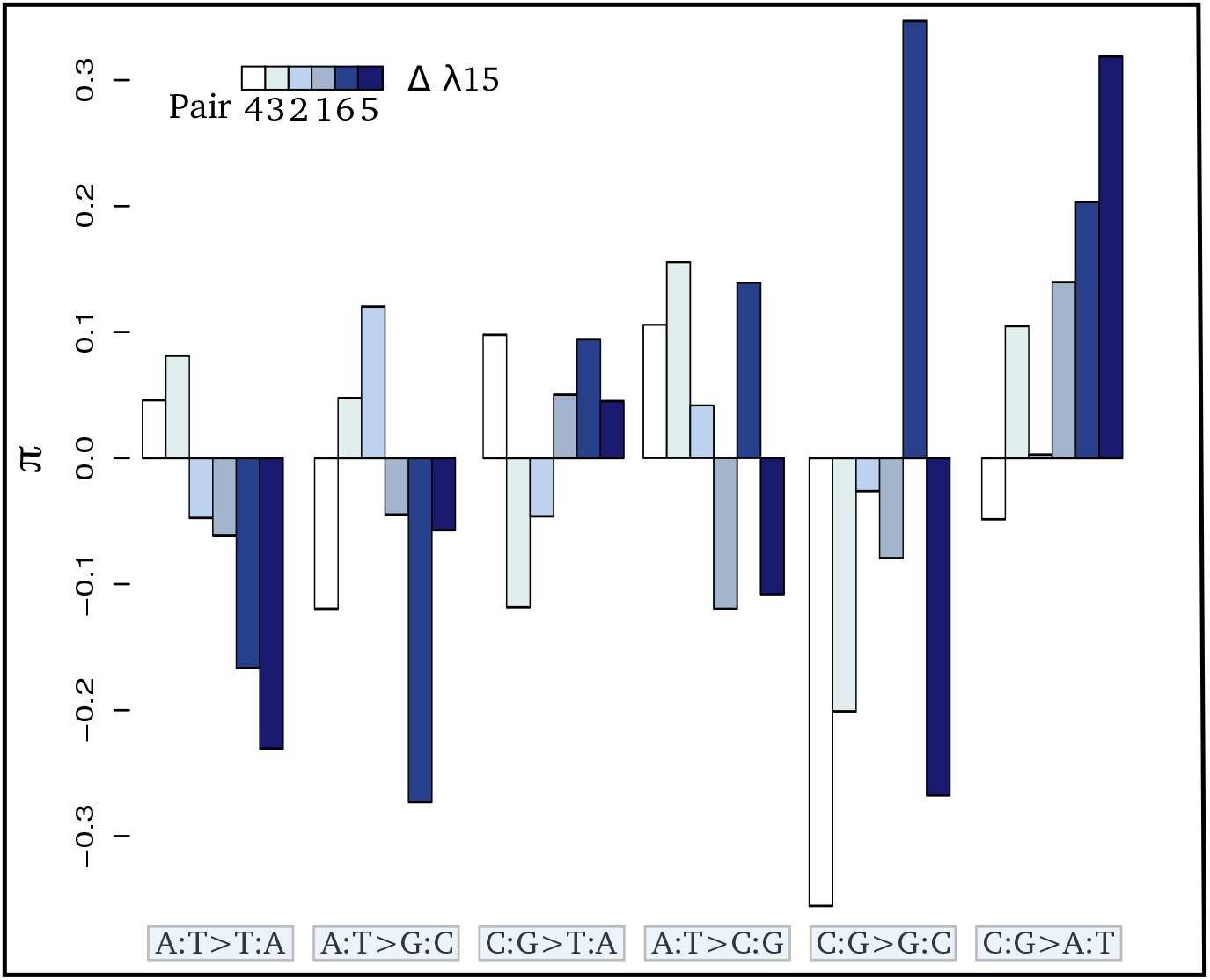
Contrasts (*π*) of the relative proportion of each mutation [*P* (*i*: *j → k*: *l*)] computed only on third positions in each pair of sister species: 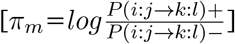 where + and - refer respectively to the species exposed to the higher and lower level of radioactivity in the pair m. Thus, positive bars represent a higher proportion of the given mutation in the species living in the high radioactivity rock. From left to right, bars are in increasing order of difference (Δ) in *λ*15 (the areal proportion of igneous and metamorphic rock in a radius of 15 km around the site) between the two species of each pair. From left to right, mutations are in increasing order of correlation with radioactivity. Numbers below the color scale indicate the species pair number as in Figure 1.

**Table S1:** 14 sequenced species with sampling coordinates, synonymous substitution rate relative to the root age for nuclear and mitochondrial genome, *α* radioactivity measured on each site, effective dose of radioactivity (in *μ*Gy/h), and this effective dose corrected for recent human impact (in *μ*Gy/h), non-synonymous substitution rate over synonymous substitution rate (*d*_*N*_ /*d*_*S*_), the areal proportion of magmatic and metamorphic rocks in a radius of 15 km around the sampling point (*λ*15), the GC content for all positions or for third positions, the effective number of codon (ENC) and the *d*_S_ computed with the CodeMl programm and with the MapNH programm.

| Species | RNA Code | $\alpha$ Radio. | lat. | long. | $d_S/\text{ra nucl}$ | $d_S/\text{ra mito.}$ | Dose |
| --- | --- | --- | --- | --- | --- | --- | --- |
| *Proasellus nsp2* MONTBAR | PWM | 0.300 | 47.01 | 5.62 | 0.1549 | 1.5924 | 1.84 |
| *Proasellus nsp* VIELVIC | PWVi | 0.900 | 44.40 | 3.94 | 0.2167 | 2.9317 | 5.29 |
| *Proasellus nsp* ROSSFELD | PWRo | 0.204 | 48.33 | 7.63 | 0.1258 | 1.3395 | 1.54 |
| *Proasellus nsp* HYPOPRAT | PWH | 1.305 | 44.01 | 3.67 | 0.2109 | 2.3386 | 7.74 |
| *Proasellus nsp* SCENOIRE | PStN | 0.480 | 47.32 | 6.43 | 0.1358 | 1.8241 | 2.22 |
| *Proasellus nsp* BRETEMIN | PStB | 2.027 | 46.20 | 4.51 | 0.1156 | 1.8590 | 33.8 |
| *Proasellus nsp1* MONTBAR | PCMt | 0.300 | 47.01 | 5.62 | 0.0481 | 1.4223 | 1.84 |
| *Proasellus nsp* GROTTAZE | PCG | 0.933 | 46.43 | 4.76 | 0.0472 | 1.6664 | 2.86 |
| *Proasellus nsp* VIONENE | PspVi | 0.350 | 44.08 | 7.10 | 0.2117 | 1.4174 | 1.53 |
| *Proasellus nsp* BOREON | ISPB | 2.370 | 44.07 | 7.25 | 0.2142 | 1.3483 | 37.2 |
| *Proasellus alibigensis* | PAL | 0.660 | 43.91 | 2.23 | 0.1042 | 2.0817 | 2.32 |
| *Proasellus nsp* GOUDECHE | PspAG | 2.413 | 44.35 | 3.78 | 0.1169 | 1.7065 | 14.7 |
| *Proasellus slavus* | slavPWRo | 0.204 | 48.33 | 7.63 | 0.1571 | 0.5893 | 1.54 |
| *Proasellus boui* | PBF | 0.748 | 44.12 | 3.89 | 0.2235 | 0.6813 | 6.49 |

| Species | $d_N/d_S$ | $\lambda_{15}$ | GC | GC3 | ENC | $d_S$ CodeML | $d_S$ MapNH | Pair |
| --- | --- | --- | --- | --- | --- | --- | --- | --- |
| *Proasellus nsp2* MONTBAR | 0.1526 | 0.1387 | 0.4041 | 0.3297 | 50.17 | 0.012413 | 0.01036720 | NA |
| *Proasellus nsp* VIELVIC | 0.1774 | 0.8985 | 0.4040 | 0.3294 | 50.18 | 0.014071 | 0.01324370 |  |
| *Proasellus nsp* ROSSFELD | 0.1503 | 0.0000 | 0.4045 | 0.3301 | 50.22 | 0.011268 | 0.01105270 | 1 |
| *Proasellus nsp* HYPOPRAT | 0.1809 | 0.6672 | 0.4041 | 0.3295 | 50.17 | 0.013782 | 0.01289470 |  |
| *Proasellus nsp* SCENOIRE | 0.1610 | 0.0000 | 0.4051 | 0.3318 | 50.35 | 0.018364 | 0.01817050 | 2 |
| *Proasellus nsp* BRETEMIN | 0.1534 | 0.6250 | 0.4051 | 0.3318 | 50.33 | 0.015513 | 0.01536320 |  |
| *Proasellus nsp1* MONTBAR | 0.1970 | 0.1387 | 0.4055 | 0.3323 | 50.36 | 0.007196 | 0.00622081 | 3 |
| *Proasellus nsp* GROTTAZE | 0.1850 | 0.1190 | 0.4054 | 0.3321 | 50.34 | 0.006391 | 0.00606323 |  |
| *Proasellus nsp* VIONENE | 0.1656 | 0.3346 | 0.4062 | 0.3347 | 50.48 | 0.016790 | 0.01658250 | 4 |
| *Proasellus nsp* BOREON | 0.1517 | 0.5306 | 0.4062 | 0.3347 | 50.46 | 0.017109 | 0.01674480 |  |
| *Proasellus alibigensis* | 0.1369 | 0.2991 | 0.3985 | 0.3163 | 49.35 | 0.029319 | 0.02895040 | 5 |
| *Proasellus nsp* GOUDECHE | 0.1369 | 0.9383 | 0.3979 | 0.3149 | 49.27 | 0.033744 | 0.03296400 |  |
| *Proasellus slavus* | 0.1672 | 0.0000 | 0.3969 | 0.3119 | 49.17 | 0.041241 | 0.04523370 | 6 |
| *Proasellus boui* | 0.1265 | 0.6645 | 0.3975 | 0.3128 | 49.19 | 0.061836 | 0.05802430 |  |

**Table S2:**
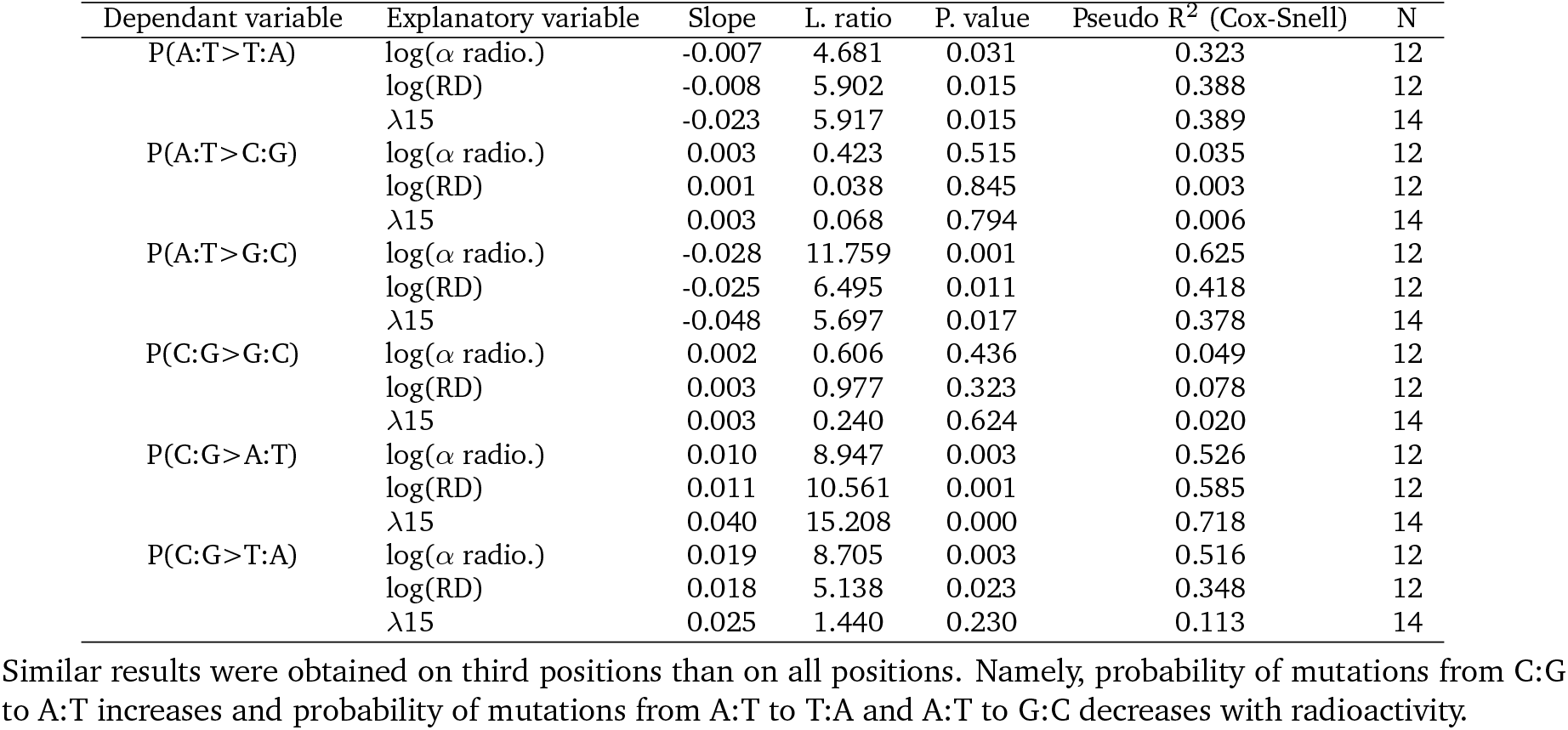
Phylogenetic Least Square (PGLS) regression of mutation probabilities computed on third positions against radioactivity measured as the *α* radioactivity measured in sediments, as the effective dose received by organisms (RD) or as the areal proportion of metamorphic and magmatic rocks in a radius of 15 km around the sampled point (*λ*15). Each line corresponds to one likelihood ratio test between the models with and without the given explanatory variable. *α* radioactivity and received dose were log transformed to fit with linear model assumptions.

| Dependant variable | Explanatory variable | Slope | L. ratio | P. value | Pseudo R<sup>2</sup> (Cox-Snell) | N |
| --- | --- | --- | --- | --- | --- | --- |
| P(A:T>T:A) | log($\alpha$ radio.) | -0.007 | 4.681 | 0.031 | 0.323 | 12 |
|  | log(RD) | -0.008 | 5.902 | 0.015 | 0.388 | 12 |
| | $\lambda 15$ | -0.023 | 5.917 | 0.015 | 0.389 | 14 |
| P(A:T>C:G) | log($\alpha$ radio.) | 0.003 | 0.423 | 0.515 | 0.035 | 12 |
|  | log(RD) | 0.001 | 0.038 | 0.845 | 0.003 | 12 |
| | $\lambda 15$ | 0.003 | 0.068 | 0.794 | 0.006 | 14 |
| P(A:T>G:C) | log($\alpha$ radio.) | -0.028 | 11.759 | 0.001 | 0.625 | 12 |
|  | log(RD) | -0.025 | 6.495 | 0.011 | 0.418 | 12 |
| | $\lambda 15$ | -0.048 | 5.697 | 0.017 | 0.378 | 14 |
| P(C:G>G:C) | log($\alpha$ radio.) | 0.002 | 0.606 | 0.436 | 0.049 | 12 |
|  | log(RD) | 0.003 | 0.977 | 0.323 | 0.078 | 12 |
| | $\lambda 15$ | 0.003 | 0.240 | 0.624 | 0.020 | 14 |
| P(C:G>A:T) | log($\alpha$ radio.) | 0.010 | 8.947 | 0.003 | 0.526 | 12 |
|  | log(RD) | 0.011 | 10.561 | 0.001 | 0.585 | 12 |
| | $\lambda 15$ | 0.040 | 15.208 | 0.000 | 0.718 | 14 |
| P(C:G>T:A) | log($\alpha$ radio.) | 0.019 | 8.705 | 0.003 | 0.516 | 12 |
|  | log(RD) | 0.018 | 5.138 | 0.023 | 0.348 | 12 |
| | $\lambda 15$ | 0.025 | 1.440 | 0.230 | 0.113 | 14 |

Similar results were obtained on third positions than on all positions. Namely, probability of mutations from C:G to A:T increases and probability of mutations from A:T to T:A and A:T to G:C decreases with radioactivity.

**Table S3:** Phylogenetic Least Square (PGLS) regression of *d*_S_ computed with CoEvol, CodeMl or mapNH against radioactivity measured as the *α* radioactivity measured in sediments, as the effective dose received by organisms or as the areal proportion of metamorphic and magmatic rocks in a radius of 15km around the sampled point (*λ*15). Each test corresponds to one likelihood ratio test between the models with and without the given explanatory variable. For *α* radioactivity and received dose, sampled sites with a break in the secular equilibrium were removed, resulting in tests with only 12 taxa.

| | $d_S$ CoEvol | | | $d_S$ CodeML | | | $d_S$ MapNH | | | N |
| --- | --- | --- | --- | --- | --- | --- | --- | --- | --- | --- |
|  | Slope | p.value | Cox-Snell R<sup>2</sup> | Slope | p.value | Cox-Snell R<sup>2</sup> | Slope | p.value | Cox-Snell R<sup>2</sup> |  |
| log($\alpha$ radio.) | 0.034 | 0.014 | 0.39 | 0.054 | 0.021 | 0.36 | 0.054 | 0.018 | 0.37 | 12 |
| log(Received Dose) | 0.038 | 0.011 | 0.42 | 0.057 | 0.024 | 0.35 | 0.057 | 0.023 | 0.35 | 12 |
| $\lambda 15$ | 0.076 | 0.003 | 0.47 | 0.118 | 0.006 | 0.42 | 0.118 | 0.005 | 0.43 | 14 |

